# A COMPARATIVE ANALAYSIS OF INTER-SITE GENE EXPRESSION HETEROGENICITY OF NORMAL HUMAN BUCCAL MUCOSA WITH NORMAL GINGIVAL MUCOSA

**DOI:** 10.1101/2021.10.08.463654

**Authors:** Thavarajah Rooban, Kannan Ranganathan

## Abstract

**BACKGROUND:** Description of heterogeneity of gene expression of various human intraoral sites are not adequate. The aim of this study was to explore the difference of gene expression profiles of whole tissue obtained from apparently normal human gingiva and buccal mucosa (HGM, HBM).

**MATERIALS AND METHODS:** Gene sets fulfilling inclusion and exclusion criteria of HGM and HBM in gene Expression Omnibus(GEO) database were identified, segregated, filtered and analysed using the ExAtlas online web tool using pre-determined cut-off. The differentially expressed genes were studied for epithelial keratinization related, housekeeping(HKG), extracellular matrix related(ECMRG) and epithelial-mesenchymal transition related genes(EMTRGs).

**RESULTS:** In all 40 HBM and 64 HGM formed the study group. In all there were 18012 significantly expressed genes. Of this, 1814 were over-expressed and 1862 under-expressed HBM genes as compared to HGM. One in five of all studied genes significantly differed between HBM and HGM. For the keratinization genes, 1 in 6 differed. One of every 5 HKG-proteomics genes differed between HBM and HGM, while this ratio was 1-in 4 for all ECMRGs and EMTRGs.

**DISCUSSION:** This difference in the gene expression between the HBM and HGM could possibly influence a multitude of biological pathways. This result could explain partly the difference in clinicopathological features of oral lesions occurring in HBM and HGM. The innate genotypic difference between the two intra-oral niches could serve as confounding factor in genotypic studies. Hence studies that compare the HBM and HGM should factor-in these findings while evaluating their results.

## 1. BACKGROUND

Gene expression serves as a bridge between information encoded within a gene and the final functional gene product, such as a protein or non-coding RNA. The orchestrated process of gene expression can be altered at any stage to modify the quantity and spatiotemporal parameters of the functional protein appearance. This is essential for preserving normal cellular structure and function. The cellular capacity to control gene expression permits it to supply a functional protein whenever it is required for its normal survival or function. This system is involved in a variety of physiological and pathological processes, including cellular adaptation to new surroundings, homeostasis maintenance, and damage recovery.^[1]^ Gene expression widely differs between the various tissues and site. It follows “tissue-specific” or signature expression patterns, reflecting possibly the differences in metabolic activity and cyto-architecture.^[2]^ Generally, all human cells contain identical housekeeping genes (the genes in every tissue; maintains cellular functions).

Fibroblasts from different parts of the human body exhibit heterogeneity with respect to cell behaviour, proliferative potential, response to growth factors and matrix biosynthesis. They also exhibit functional specialization according to the tissue of origin, site, and spatial location, even in the same tissues. Intra-site and inter-site heterogeneity of fibroblasts have been characterized in tissues like gingiva, periodontal ligament, and forearm skin. They often differ in their proliferative potential, ability to form colonies in semi-solid medium, cellular response to cytokines, secretion of matrix-degrading enzymes, migratory behaviour, cytokine production and matrix deposition.^[3]^

Fibroblast of anatomically distinct sites have distinct transcriptional patterns. On appropriate stimulation, the relatively quiescent fibroblasts can acquire an active synthetic, contractile phenotype and express several smooth muscle cell markers, which are not exclusive for fibroblasts of that particular niche.^[4,5]^ Failure to account and consider the phenotypic as well as expression profile of these heterogenic differences can lead to potential erroneous interpretation of data collected for the specific experimental purposes.^[6]^ All oral fibroblasts are derived from the ecto-mesenchymal cells of the cranially migrated neural crest cells. In spite of the same lineage, there is anatomical, histological heterogeneity among various intra-oral sites.^[7]^

Human buccal fibroblasts (HBF) have partly primed, committed cells of neural crest lineage characterised by plasticity and longevity controlled by WNT and p53 gene network.^[8]^ The existence of heterogeneity in human gingival fibroblasts (HGF) is documented, particularly those of the gingival papillary and reticular area.^[9,10]^ A subset population of HGF is known to heal without fibrosis.^[11]^ When HGF and HBF cultured fibroblasts were studied, the cell density migration index was higher in HBF than HGF. On the other hand, the HGF had more adult phenotype in contrast to foetal phenotype of HBF.^[9]^

Oral epithelial cells are exposed to external environment and they also interact extensively with underlying cells such as fibroblasts and immune cells. Alteration in fibroblasts influences proliferation, repair and inflammatory cytokine secretion and factors released by fibroblasts influence overlying epithelial cells.^[12]^ Oral keratinocytes interact with themselves and underlying connective tissue to provide a tight barrier and heal when damaged.^[13-16]^

The aim of this manuscript was to study the difference of expression profiles of epithelial and the extracellular matrix (ECM) components of whole tissue obtained from two common oral niches – gingiva and buccal mucosa from publicly available Gene Expression Omnibus - GEO datasets.

## 2. MATERIALS AND METHODS

### 2.1 Source of microarray datasets

A systematic search of the National Centre for Biotechnology Information (NCBI-GEO - http://www.ncbi.nlm.nih.gov/geo) repository using the keywords “buccal mucosa” “gingiva” was carried out. The microarray datasets and their descriptions contents were further limited to a. in Humans - *Homo sapiens* b. type - “expression profiling by the array.”, c. has normal healthy patient, tissue biopsy (epithelium and connective tissue) sample with no obvious disease d. ideal, acceptable extraction protocol e. has mRNA. Datasets that were from culture explants or had only fibroblasts or only keratinocytes or non-tissues or that unclear extraction protocols or sources or details were excluded. No emphasis was placed on types of dataset platforms. The gene series were collated.

### 2.2 Pair-Wise Comparison

The individual patient’s gene expression datasets from methodically reviewing and screening the microarray datasets were collated using the pair wise comparison using the ExAtlas online web tool at https://lgsun.irp.nia.nih.gov/exatlas.^[17]^ The genes from the GEO datasets were log_2_ transformed, normalized with quantile normalization method and later combined. Then, the samples – Human Buccal Mucosa (HBM) and Human Gingival Mucosa (HGM) gene-sets were combined. Principal component analysis was performed to check the tissue/gene distribution with FDR ≤0.05, correlation threshold = 0.7 and fold change threshold of 2. In the tool, PCA is calculated using the Singular Value Decomposition (SVD) method that generates eigenvectors for rows as well as columns of the log-transformed data matrix.^[17]^ For plotting of tissues and genes (biplot) column projections were used as biplots is helpful for visually exploring associations between genes and tissues.

A pairwise compared with ANOVA statistical methodology was employed to compare HBM and HGM. The quality assessment measurement of the samples within the pooled datasets was the standard deviation (SD) value, and the criterion was SD ≤ 0.3. The correlation of gene expression for housekeeping genes was established at >0.5. The false discovery rate (FDR) based on the Benjamini-Hochberg was ≤0.05 and fold change was fixed at 2. The scatter plot was made to identify the over-expressed and under-expressed genes in HBM as compared to HGF. Only genes that had gene symbol was considered for further studies.

### 2.3 Gene-set enrichment analysis of up/down-regulated genes

The listed genes were then used to compare with the Public human Gene Ontology gene set with functional role of 9606 genes.^[18,19]^ A FDR≤0.05, fold enrichment threshold of 2 and a minimum of 5 gene overlap in biological process was kept as standard norms. For this purpose, exatlas tool employs, Parametric Analysis of Gene Enrichment (PAGE) as it is simple and reliable. ^[20,21]^

### 2.4 Differential expression of epithelial keratinization related genes

The genes associated with epithelial keratinization were collected from gene ontology (GO: 0031424; http://amigo.geneontology.org/amigo/term/GO:0031424). The differentially expressed genes from section-2.2 were assessed for the same.

### 2.5 Differential expression of housekeeping gene

From a database of human housekeeping genes (HKG-proteomics) (https://www.proteinatlas.org/humanproteome/tissue/housekeeping), the results of section-2.2 were compared and the housekeeping genes that were differentially expressed between HBM and HGM tabulated.^[22]^

### 2.6 Differential expression of extracellular matrix related genes [ECMRGs]

The list of over and under-expressed genes of HBM as compared to HGM was searched for extracellular matrix (ECM) and annotated according to matrisome divisions (core matrisome or matrisome-associated) and categories (ECM glycoproteins, collagens and proteoglycans, for core matrisome genes, ECM-affiliated, ECM regulators and secreted factors, for matrisome-associated genes) in lines as proposed by Naba et al., using the MatrisomeAnnotator tool at http://matrisomeproject.mit.edu/analytical-tools/matrisome-annotator/^[23]^

### 2.7 Differential expression of epithelial-mesenchymal transition related genes [EMTRGs]

From the database of human epithelial-mesenchymal transition (EMT) database, the results from section-2.2 were compared and the differentially expressed EMT genes were listed out.^[24]^

### 2.8 Differential expression of fibroblast markers

The list of fibroblast markers of heterogeneity as revealed by recent single-cell analysis through cell identification and discrimination was collected from previously published literature.^[25]^ A comparison of the same was performed based on ANOVA results and the overlap tabulated along with rank.

## 3. RESULTS

### 3.1 Microarray datasets

From the publicly available database, in early March 2021, a total of 4 datasets (GSE7307, n=4; GSE3526, n=4; GSE17913, n=40) non-smoker HBM and 5 datasets (GSE106090, n=6; GSE10334, n=64; GSE4250, n=2; GSE3374, n=8; GSE23586, n=3) had healthy HGM were collected and processed in the Exatlas software as outlined. In all there were 48 HBM and 83 HGM gene-sets. All these gene-sets were subjected to quality assessment procedure as previously outlined. Those sets whose house keeping genes did not fulfil the required minimum quality parameters of correlation or SD were removed from further analysis. After removal of non-qualifying datasets, a total of 40 HBM and 64 HGM formed the final study group. The final study group gene-set had used GPL570 platform to analyse the array.

### 3.2. Pair-wise comparison

The final combined dataset had 104 samples comprising 22738 probes. Of this, 22139 had gene symbol. Of this there were 18012 genes with FDR≤ 0.05 and included in the PCA. The relationship of tissues and genes are shown in figure-1A, 1B. Of the PCA, the component 1 had an Eigen value of 27.17 accounting for 90.16% of all genes (Figure-2). The scatter-plot was obtained (Figure-3) with HBM showing 1843 (1814 with gene symbols) overexpressed genes and 1915 (1862 with gene symbols) under-expressed genes as compared to HGM. In all, of the 18012 genes, 3758(20.86%) were differentially expressed with statistical significance.

**FIGURE 1:**
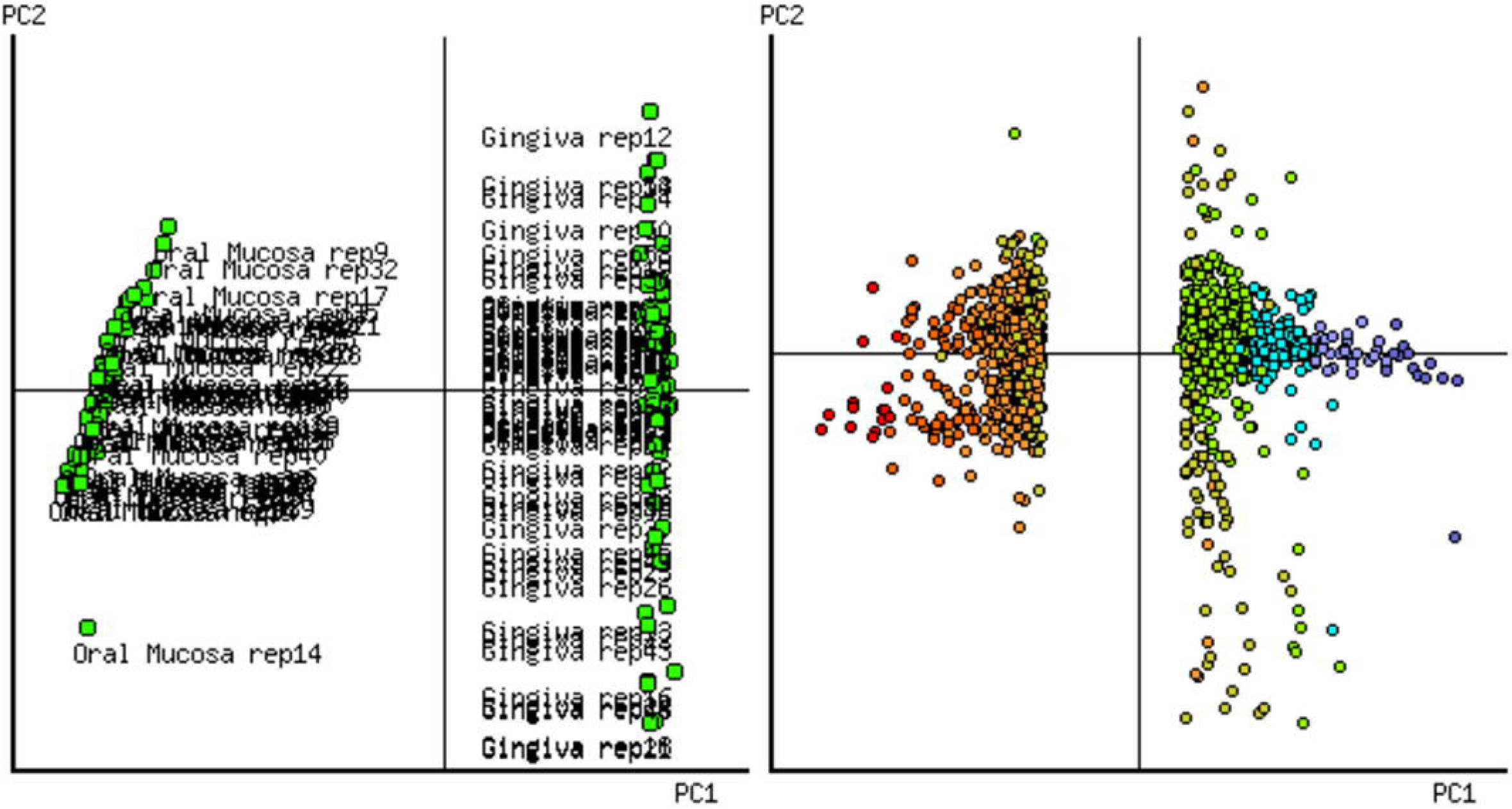
Relationship between the Human Buccal Mucosa and Human Gingival Mucosal Samples A. Tissue B. Genes in Principal Component Analysis

**FIGURE-2:**
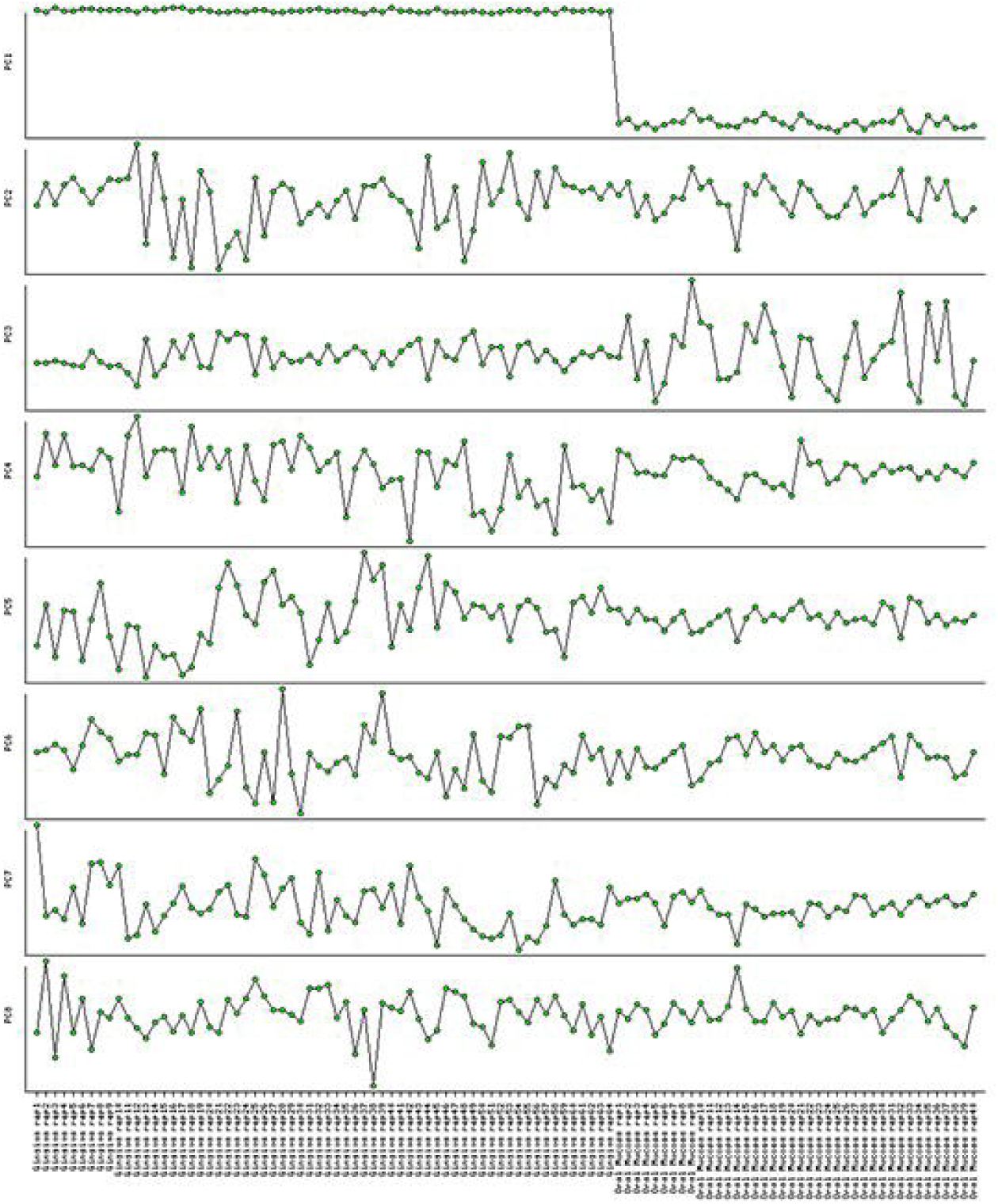
Different components of the Principal Component Analysis of various samples studied.

**FIGURE-3:**
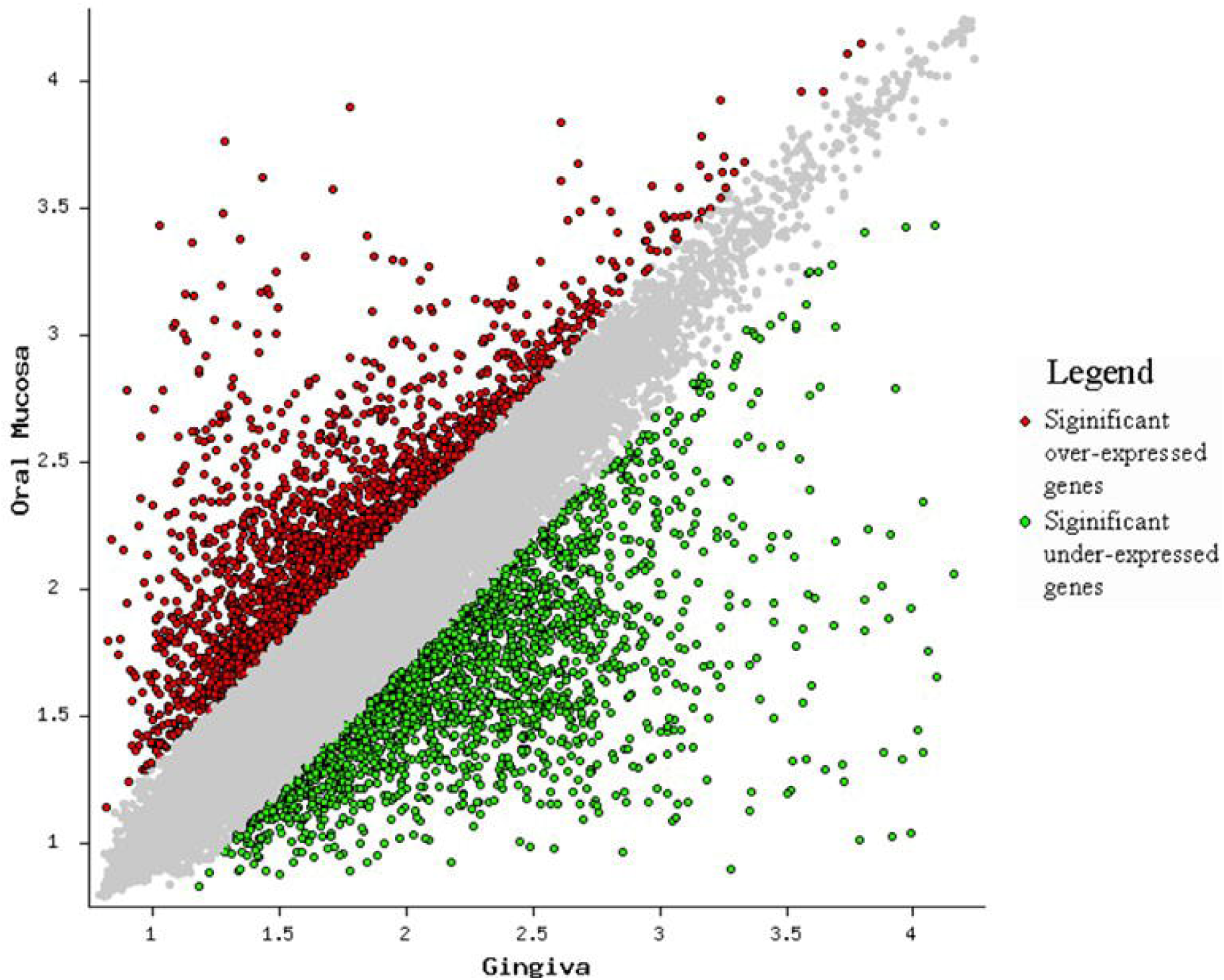
Scatter-plot showing the differentially expressed genes between Human Buccal Mucosal and Human Gingival Mucosal Samples.

**FIGURE-4:**
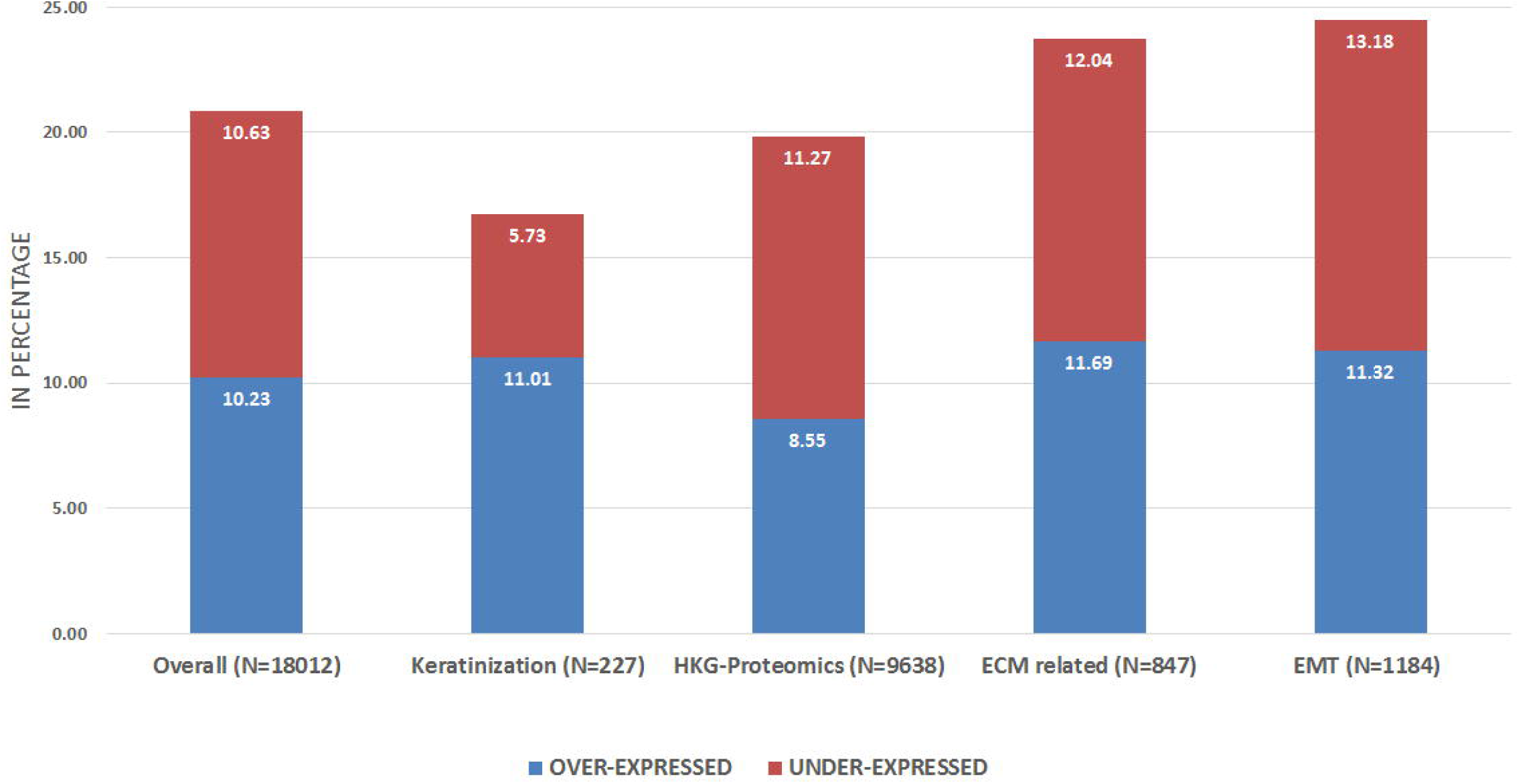
Overall differences between various gene expressions of Human Buccal Mucosa and Human Gingival Mucosal Samples.

### 3.3 Gene-set enrichment analysis of up/down-regulated genes

The over/under expressed gene enrichment analysis using Gene-Ontology is shown in supplementary file-1, (Tables1, 2). There were 217 biological processes in over-expressed genes. The commonly involved GO processes (>15% of cluster) and their cluster frequency included anatomical structure development (GO:0048856, 37.14%), biosynthetic process(GO:0009058, 33.64%), cellular nitrogen compound metabolic process(GO:0034641, 31.34%), signal transduction(GO:0007165, 30.62%), transport(GO:0006810, 28.2%), cell differentiation(GO:0030154, 26.39%), cellular protein modification process(GO:0006464, 21.86%), response to stress(GO:0006950, 21.86%), cellular component assembly(GO:0022607, 17.69%) and catabolic process (GO:0009056, 16.3%).

**Table-1:**
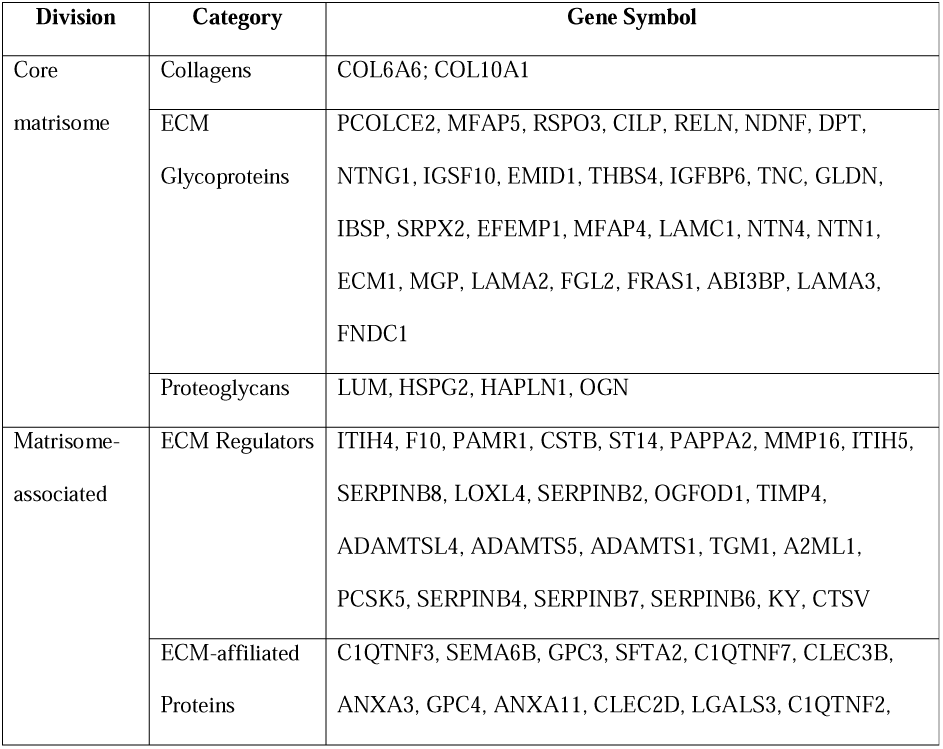
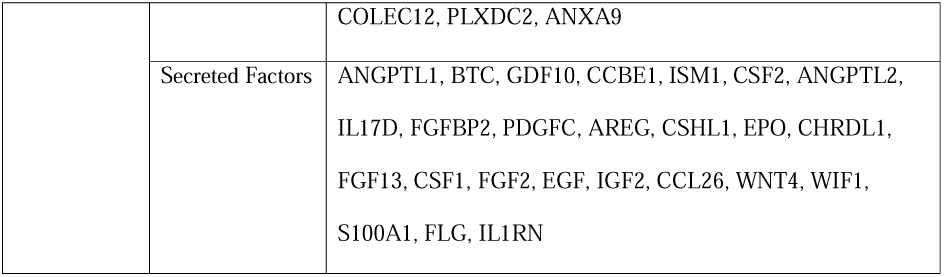
Matrisome related protein’s mRNA over expressed in Human Buccal mucosa (n=40) as compared to Human Gingival mucosa (n=64)

**Table-2:** Matrisome related protein’s mRNA under expressed in Human Buccal mucosa (n=40) as compared to Human Gingival mucosa (n=64)

| Division | Category | Gene Symbol |
| --- | --- | --- |
| Core matrisome | Collagens | COL11A1, COL13A1, COL4A5, COL5A2, COL6A3, COL3A1, COL8A1, COL4A4, COL27A1, COL12A1, COL4A1, COL4A2 |
|  | ECM Glycoproteins | HMCN2, EDIL3, LAMB3, CTHRC1, CRISPLD1, THBS1, AGRN, THBS2, SPARC, POSTN, CRISPLD2, NID2, PCOLCE, SPP1, VWA3A, TNFAIP6 |
|  | Proteoglycans | ASPN, SRGN |
| Matrisome-associated | ECM Regulators | CST3, CD109, ADAMTS2, LOX, SERPINB13, MMP11, ADAMDEC1, TGM7, MMP15, ADAM22, EGLN2, PRSS12, MMP12, HYAL1, MMP9, MMP19, ADAMTS6, NGLY1, ADAM28, MASP1, MMP7, EGLN3, MMP13, SERPINA1, TIMP2, LOXL2, BMP1, ADAM12, MMP10, MMP1, SERPINB5, MMP3, FAM20A, PCSK6, FAM20C, SERPINE1, PI3 |
|  | ECM-affiliated Proteins | ANXA2, CLEC4E, CLEC7A, C1QTNF1, SEMA5B |
|  | Secreted Factors | IL22, MEGF9, WNT5A, TNFSF10, CXCL13, RPTN, CXCL1, S100A7, INHBA, CBLN2, CCL28, CXCL8, CXCL2, CCL20, IL19, CCL4, CXCL6, CCL3, CXCL3, CXCL5, BMP6, IL24, FRZB, S100A7A, CCL2, IL6, FLG2, CCL19, SFRP1, CXCL10 |

The under-expressed genes were associated with 192 biological processes. Of the under-expressed processes in HBM, the most common include ECM, CXCR chemokine, hemidesmosomes, collagen-activated tyrosine kinase receptor signalling pathway as compared to HGM. (Supplemental file-1, tables-3,4) Of the under-expressed genes, the commonly involved GO processes (>15% of cluster) and their cluster frequency included cellular nitrogen compound metabolic process (GO:0034641; 36.81%), anatomical structure development (GO:0048856; 35.17%), biosynthetic process (GO:0009058; 34.58%), signal transduction (GO:0007165; 34.23%), transport (GO:0006810; 32.24%), cellular protein modification process (GO:0006464; 25.73%), cell differentiation (GO:0030154; 25.5%), response to stress (GO:0006950; 24.97%), immune system process (GO:0002376; 20.75%), catabolic process (GO:0009056; 19.17%), cellular component assembly (GO:0022607; 17.53%) and cell death (GO:0008219; 15.01%).

**Table-3:** Expression of fibroblastic markers in the Human Buccal mucosa (n=40) as compared to Human Gingival mucosa (n=64)

| **Gene Symbol** | **Average Intensity** | **Human buccal Mucosa (n =40)** | **Human Gingival mucosa (n=64)** | **Variance (Factor)** | **Variance (Err)** | **Smooth Variance (Err)** | **Final MSE** | **F-statistics** | **P** | **FDR** | **rank** |
| --- | --- | --- | --- | --- | --- | --- | --- | --- | --- | --- | --- |
| PCOLCE2 | 1.7571 | 2.418 | 1.0961 | 43.01233 | 0.0165 | 0.01517 | 0.0165 | 2606.325 | 0 | 0 | 339 |
| MFAP5 | 1.7253 | 2.2919 | 1.1587 | 31.61427 | 0.02545 | 0.01496 | 0.02545 | 1242.374 | 0 | 0 | 698 |
| ADAMTS2 | 2.0129 | 1.5902 | 2.4356 | 17.59442 | 0.02881 | 0.01644 | 0.02881 | 610.774 | 0 | 0 | 1325 |
| LUM | 2.5967 | 3.0943 | 2.0992 | 24.37275 | 0.04575 | 0.01431 | 0.04575 | 532.724 | 0 | 0 | 1503 |
| CELF2 | 1.8467 | 2.2209 | 1.4726 | 13.78332 | 0.02733 | 0.01635 | 0.02733 | 504.317 | 0 | 0 | 1574 |
| DPT | 2.3227 | 2.7092 | 1.9362 | 14.70734 | 0.03152 | 0.01554 | 0.03152 | 466.55 | 0 | 0 | 1686 |
| S100A16 | 3.9037 | 3.9999 | 3.8075 | 0.91167 | 0.00266 | 0.00906 | 0.00266 | 343.166 | 0 | 0 | 2229 |
| IGFBP6 | 3.1453 | 3.4865 | 2.804 | 11.46702 | 0.03519 | 0.01175 | 0.03519 | 325.84 | 0 | 0 | 2319 |
| NAV1 | 2.0801 | 1.864 | 2.2962 | 4.59665 | 0.01755 | 0.01614 | 0.01755 | 261.848 | 0 | 0 | 2868 |
| MFAP4 | 2.357 | 2.6097 | 2.1042 | 6.29119 | 0.02465 | 0.01513 | 0.02465 | 255.242 | 0 | 0 | 2927 |
| ABCA8 | 2.3259 | 2.7553 | 1.8966 | 18.14898 | 0.072 | 0.01562 | 0.072 | 252.071 | 0 | 0 | 2949 |
| LPAR1 | 2.2663 | 2.4833 | 2.0493 | 4.63588 | 0.02162 | 0.01644 | 0.02162 | 214.441 | 0 | 0 | 3393 |
| SCARA5 | 2.4022 | 2.7477 | 2.0568 | 11.75022 | 0.063 | 0.01511 | 0.063 | 186.526 | 0 | 0 | 3777 |
| RNASE4 | 2.2312 | 2.3937 | 2.0686 | 2.60063 | 0.0142 | 0.01681 | 0.0142 | 183.19 | 0 | 0 | 3844 |
| GSN | 2.4363 | 2.6528 | 2.2198 | 4.61547 | 0.02754 | 0.01483 | 0.02754 | 167.571 | 0 | 0 | 4160 |
| MEDAG | 1.9843 | 2.1431 | 1.8255 | 2.48321 | 0.01925 | 0.01573 | 0.01925 | 128.986 | 0 | 0 | 5100 |
| PI16 | 1.9328 | 2.0479 | 1.8177 | 1.30357 | 0.01135 | 0.01683 | 0.01135 | 114.884 | 0 | 0 | 5579 |
| HTRA3 | 1.7017 | 1.7924 | 1.611 | 0.8096 | 0.00849 | 0.01516 | 0.00849 | 95.385 | 0 | 0 | 6319 |
| DCN | 3.4857 | 3.3553 | 3.6161 | 1.67483 | 0.02221 | 0.0101 | 0.02221 | 75.395 | 0 | 0 | 7334 |
| CD34 | 2.0667 | 2.1721 | 1.9614 | 1.09282 | 0.01477 | 0.01654 | 0.01477 | 73.979 | 0 | 0 | 7428 |
| C3 | 2.5953 | 2.768 | 2.4226 | 2.93593 | 0.04736 | 0.01433 | 0.04736 | 61.986 | 0 | 0 | 8222 |
| HSD11B1 | 1.5493 | 1.4199 | 1.6787 | 1.6479 | 0.02792 | 0.01516 | 0.02792 | 59.025 | 0 | 0 | 8465 |
| FBLN1 | 2.5693 | 2.7181 | 2.4204 | 2.18212 | 0.03812 | 0.01379 | 0.03812 | 57.241 | 0 | 0 | 8647 |
| LSP1 | 2.2338 | 2.3348 | 2.1328 | 1.00462 | 0.02097 | 0.0167 | 0.02097 | 47.897 | 0 | 0 | 9447 |
| DPEP1 | 1.7591 | 1.7098 | 1.8085 | 0.24006 | 0.00634 | 0.015 | 0.00634 | 37.867 | 0 | 0 | 10539 |
| COL5A1 | 2.3418 | 2.2464 | 2.4371 | 0.89482 | 0.03278 | 0.01515 | 0.03278 | 27.299 | 0 | 0 | 12004 |
| FBLN2 | 2.2682 | 2.1947 | 2.3418 | 0.53318 | 0.02114 | 0.01631 | 0.02114 | 25.218 | 0 | 0 | 12322 |
| COL1A2 | 2.9908 | 2.9026 | 3.079 | 0.7663 | 0.0378 | 0.01189 | 0.0378 | 20.271 | 0.00002 | 0.00003 | 13194 |
| LOXL1 | 2.176 | 2.1031 | 2.2489 | 0.52319 | 0.02654 | 0.01643 | 0.02654 | 19.713 | 0.00002 | 0.00004 | 13309 |
| MMP2 | 2.7886 | 2.6945 | 2.8827 | 0.87235 | 0.04547 | 0.01264 | 0.04547 | 19.183 | 0.00003 | 0.00005 | 13405 |
| COL1A1 | 2.779 | 2.6669 | 2.8911 | 1.23757 | 0.07838 | 0.01285 | 0.07838 | 15.788 | 0.00013 | 0.00021 | 14119 |
| HEG1 | 2.4937 | 2.4358 | 2.5515 | 0.32943 | 0.02104 | 0.01392 | 0.02104 | 15.659 | 0.00014 | 0.00023 | 14148 |
| SLC43A3 | 2.1587 | 2.105 | 2.2124 | 0.28393 | 0.02048 | 0.01622 | 0.02048 | 13.861 | 0.00032 | 0.0005 | 14568 |
| CCDC80 | 2.1613 | 2.2333 | 2.0892 | 0.51087 | 0.04333 | 0.01643 | 0.04333 | 11.789 | 0.00086 | 0.0013 | 15136 |
| OLFML3 | 2.3478 | 2.408 | 2.2875 | 0.35735 | 0.03091 | 0.01531 | 0.03091 | 11.562 | 0.00096 | 0.00144 | 15189 |
| BICC1 | 1.907 | 1.8595 | 1.9544 | 0.2214 | 0.02466 | 0.01695 | 0.02466 | 8.979 | 0.00343 | 0.00486 | 16030 |
| PDGFRA | 2.8541 | 2.8137 | 2.8945 | 0.16057 | 0.02236 | 0.01219 | 0.02236 | 7.182 | 0.00859 | 0.01168 | 16717 |
| GFPT2 | 1.7615 | 1.738 | 1.785 | 0.05438 | 0.01748 | 0.0152 | 0.01748 | 3.111 | 0.08077 | 0.0981 | 18721 |
| ENTPD2 | 1.5199 | 1.5357 | 1.5041 | 0.02465 | 0.00983 | 0.01516 | 0.00983 | 2.507 | 0.11645 | 0.13831 | 19143 |
| SERPINF1 | 2.9651 | 2.9421 | 2.988 | 0.05187 | 0.02608 | 0.01205 | 0.02608 | 1.989 | 0.16153 | 0.18836 | 19499 |
| MGST1 | 2.4282 | 2.3839 | 2.4724 | 0.19169 | 0.10107 | 0.01527 | 0.10107 | 1.897 | 0.17151 | 0.19927 | 19567 |
| S100A10 | 2.6527 | 2.6331 | 2.6723 | 0.03777 | 0.0334 | 0.01415 | 0.0334 | 1.131 | 0.29006 | 0.32545 | 20265 |
| UGDH | 2.5123 | 2.501 | 2.5235 | 0.01247 | 0.01519 | 0.01382 | 0.01519 | 0.821 | 1 | 1 | 20631 |

### 3.4 Differential expression of epithelial keratinization related genes

In all there were 227 genes associated with epithelial keratinization. Of this, there were 13 genes overexpressed and 25 under-expressed in HBM as compared to HGM. Combined, this 38 genes accounted to 16.74% of all keratinization associated genes.

Of the 13 genes overexpressed in HBM, 5 were keratin related (KRT31, KRT33A, KRT33B, KRT7, KRT78). The other genes are FLG, KLK5, KRTAP3-2, PPL, PRSS8, SPINK5, ST14 and TGM1. Of the 25 under-expressed genes, 7 were keratin related (KRT2, KRT3, KRT4, KRT10, KRT16, KRT24, KRT76). The rest includes ABCA12, CAPN1, CDH3, CDSN, CYP26B1, DSC1, JUP, KAZN, LCE1B, LCE1E, LCE2B, LCE3D, LORICRIN, PCSK6, PI3, PKP2, RPTN and SPRR2G.

### 3.5 Differential expression of housekeeping gene

There were 9638 genes identified in the HKG-Proteomics database. Of this, 1910 (19.38%) of all HKG-Proteomics) were identified in the differentially expressed genes of this study. This 1910 contributed to 52% of all differentially expressed genes in this study. Of the 1910 genes, 824 (43.14%) were over-expressed and 1086 (56.86%) were under-expressed.

### 3.6 Differential expression of ECMRGs

There are 847 ECMRGs identified and reported.^[26]^ The identified over/under-expressed genes were subjected to matrisome analysis. Of the 1843 genes over-expressed in HBM as compared to HGM, there were 99 matrisome related factors (core matrisome (2 collagens - Col6A6; Col10A1; 29 ECM Glycoproteins and 4 proteoglycans), matrisome associated (24 ECM regulators, 15 ECM affiliated proteins, 25 secreted factors) (Table-1). Of the 1915 genes under-expressed in HBM as compared to HGM, there were 102 matrisome related factors (core matrisome (12 collagens; 16 ECM Glycoproteins and 2 proteoglycans), matrisome associated (37 ECM regulators, 5 ECM affiliated proteins, 30 secreted factors). (Table-2) In all, 201 ECMRGs of the 847 were differentially expressed in the present study. This accounts to 23.73% of the ECM genes.

### 3.7 Differential expression of EMTRGs

There were 1184 human EMTRGs. Of the 1843 overexpressed HBM genes, 134(7.3%) were EMT genes and 1915 under-expressed genes, 156 (8.1%) were EMT genes. This difference was not statistically significant. (P=0.173) The differentially expressed 290 EMT related genes in present study accounted to 24.49% of all known EMT genes.

### 3.8 Differential expression of fibroblast markers

Of the 45 listed markers, there were 37 distinct markers differentially expressed between HBM and HGM. Of the 37 markers, 22 genes (PCOLCE2, MFAP5, LUM, ABCA8, DPT, CELF2, SCARA5, IGFBP6, MFAP4, LPAR1, GSN, C3, RNASE4, MEDAG, FBLN1, PI16, CD34, LSP1, S100A16, HTRA3, CCDC80, OLFML3) were over-expressed in HBM while 15 genes (PDGFRA, BICC1, DPEP1, SLC43A3, HEG1, LOXL1, FBLN2, COL1A2, MMP2, COL5A1, COL1A1, HSD11B1, DCN, AV1, ADAMTS2) were under-expressed in HBM as compared to HGM. The extent of difference is listed in as table-3 with rank and desired statistics.

### 3.9 Overall difference between gene expression of HBM and HGM

Among all the significant genes in this study, 1-in-5 significantly differed between HBM and HGM. For the keratinization genes, 1 in 6 differed. One of every 5 HKG-proteomics genes differed between HBM and HGM, while this ratio was 1-in 4 for all ECMRGs and EMTRGs. (Figure-XX)

## DISCUSSION

Several studies have reported the spatio-temporal gene expression difference among various human tissues, organs and even different parts of same organ, including skin, adipose tissue, brain, cornea and epidymis.^[27-32]^ The role of heterogeneity of gene expression of fibroblasts of various sites and its implication on human diseases have been reported.^[33]^ Studies exploring the heterogeneity of the various intra-oral sites in terms of pH, histo-cyto-architecture, immune-topology and gene expressions are limited.^[7-16, 34,35]^ Existing studies have reflected the inherent differences between the various intra-oral sites and even among the same site such as gingiva and periodontium.^[9,10,11]^ The spatio-temporal heterogeneity of the gingiva at single cell level has been recently described widening our understanding of the biology of these intra-oral sites.^[36]^ The intra-oral site specific gene expression differences in pathologic process also has been previously reported.^[36,37]^ However, the innate difference between the gene expression of whole HBM and HGM among normal, non-diseased population has not been reported. Hence this study was attempted.

The present study reveals the existence of substantial difference between the HBM and HGM in terms of gene expression. The figure-xx shows the extent of difference. The difference emanates from the basic housekeeping genes, keratinization process to complex ECMRGs and EMTRGs. This inherent difference in gene expression may have ramification and may partially account for site predilection of pathologies. ^[37,38]^ In addition to differential expression, the net difference in key biological reactions between HBM and HGM have been evidenced by change in the KEGG pathways. The alterations could be reflecting on the molecular profile of the cells of HBM and HGM.

For a typical biomedical research involving molecular techniques, the need for positive and negative control is mandatory. There are guidelines for the use of such procedure related control tissues and mandatory reporting format.^[39]^ However, to the best of our knowledge, there are only few suggestions and deliberations for “normal” tissues that are involved as an experimental control arm, especially in oncology.^[40]^ Ethically, ideal normal tissues are difficult to acquire.^[41-43]^ Most of the bioethical guidelines in force advises against incising/excising or enlarging to non-lesional areas, for the exclusive purpose of conducting a research.^[42]^ Generally a normal tissue that is trimmed/excised for approximation after unrelated surgical treatment, after voluntary consent is used as normal tissues.^[43]^ This “uninvolved matching” tissue may not a true representation of a native, normal tissue because molecular changes may have occurred under the influence of adjacent infection, inflammation or an adjacent neoplastic process (“condemned mucosa”). When “normal” tissue is required as a control arm, it becomes pertinent to specify what tissues constitute “normal” and justify that such a tissue (if from diseased entity) would not have altered molecular signals that could adversely influence the outcome of the comparison.^[41]^ The influence of age, gender and habits need to be also accounted. The apparent clinically normal tissues could possibly harbour mutations and molecular changes that could potentially influence the outcome of comparisons.^[40]^ Hence the apparent healthy tissues should be used with caution.

Some investigators request normal tissues from trauma cases or edges of chronic infection and even from preserved anatomic entities. Wound approximation edges from trauma cases may be contaminated and if late, be affected by inflammatory process. Similarly chronic inflammation could alter the molecular nature of the native tissues. It may not be ethical to obtain tissues from autopsies or stored specimen because these may be done without consent.^[41]^

There has been attempts to elucidate the normal human differential protein expression in various tissues. Even in the study, different intra-oral sites have not been evaluated in such attempts except for tongue.^[44,45]^ Cancer process per se, are known to down-regulate tissue specific genes and are directly associated with prognosis. While studying differences between same type of cancer (for example oral squamous cell carcinoma) at different sites (of oral cavity), the inherent difference between the normal tissues would be highlighted more rather than differences between those that are independent of normal tissue physiology.^[46]^ If not accounted, this phenomenon may lead to erroneous conclusions.

In studies involving human oral diseases, the normal, non-diseased control tissues are often obtained from gingival tissues or from the retromolar tissue- an area where HBM and HGM meet. In such instance, with chronic exposure to pro-inflammatory and pro-fibrotic cytokines, as in inflammation, the connective tissue and epithelium may be irreversibly damaged. As a part of reparative mechanism, there could a cascade of triggered epigenetic modifications and activation of related genes, leading to their engagement in further differentiation and fibrosis development. Based on the type of cells, the reaction could also widely vary. There are gaps in knowledge regarding this complex mechanism at the cellular and molecular levels.^[47]^ The biological niches in the HBM and HGM are different. In addition, the normal tissue resident microbial flora could influence the gene expression. There are recent reports emanating from cancer and normal tissues.^[48,49]^ In such a background, the inherent geno-typological difference between HBM and HGM, as evidenced in this study assumes an important role.

Limitation of the study includes using samples from different studies and population, though the difference have been accounted for in the analysis; the samples differing from several studies but had used one platform; stringent quality control parameter being enforced; non-consideration of age, gender, and deleterious habits. Future studies need to account for the same.

## CONCLUSION

An attempt is made to characterize the genetic expression of HGM and HBM using pre-existing dataset and employing stringent statistical approach. The study identified that 1- in-5 genes significantly differed between HBM and HGM. Of genes responsible for keratinization process, 1 in 6 differed. Similarly housekeeping genes, genes associated with extracellular matrix and epithelial-mesenchymal transition were also significantly altered. This inherent genotypic difference between the 2 intra-oral niches could mislead the genotypic studies, if the differences are not properly accounted. Future studies that compare the HBM and HGM should factor-in the findings while evaluating their results. Large scale prospective studies are needed further to validate the findings.

